# Obesity impairs resistance to *Leishmania major* infection in C57BL/6 mice

**DOI:** 10.1101/342642

**Authors:** Franciele Carolina Silva, Vinicius Dantas Martins, Felipe Caixeta, Matheus Batista Carneiro, Graziele Ribeiro Goes, Nivea Carolina Paiva, Cláudia Martins Carneiro, Leda Quercia Vieira, Ana Maria Caetano Faria, Tatiani Uceli Maioli

**Author notes:** Corresponding author (MAIOLI, T.U.). These authors contribute equally (Franciele Carolina Silva, Vinicius Dantas Martins).

## Abstract

An association between increased susceptibility to infectious diseases and obesity has been described as a result of impaired immunity in obese individuals. It is not clear whether a similar linkage can be drawn between obesity and parasitic diseases. To evaluate the effect of obesity in the immune response to cutaneous *L. major* infection, we studied the ability of C57BL/6 mice submitted to a high fat and sugar diet to control leishmaniasis. Mice with diet-induced obesity presented thicker lesions with higher parasite burden and more inflammatory infiltrate in the infected ear when infected with *L. major*. We observe no difference in IFN-γ or IL-4 production by draining lymph node cells between control and obese mice, but obese mice presented higher production of IgG1 and IL-17. A higher percentage of *in vitro-*infected peritoneal macrophages was found when these cells were obtained from obese mice when compared to lean mice. *In vitro* stimulation of macrophages with IL-17 decreased the capacity of cells from control mice to kill the parasite. Moreover, macrophages from obese mice presented higher arginase activity. Together our results indicate that diet-induced obesity impairs resistance to *L. major* in C57BL/6 mice without affecting the development of Th1 response.

**Author Summary:** The obesity is a public health problem and it is reaching extraordinary numbers in the world and others diseases are being involved and aggravated as consequence of obesity. What we know is that some diseases are more severe in obese people than in normal people. We did not know how obesity changes the profile of immune response to infectious agents, leading to the more severe diseases. That‘s why we decided to investigate how obese mice lead with *Leishmania major* infection. Leishmaniasis is a protozoa parasite infection considered a neglected disease. To try our hypothesis we gave a hipercaloric diet to induce obesity in C57BL/6 mice. After that, we injected *L. major* in the mice ear and followed the lesion for 8 weeks. We observed a ticker lesion and the cells from draining lymph node from obese mice produced more IL-17 than cells from normal mice. We also infected in *vitro*, macrophages from obese mice and stimulated the cells with IL-17, and we observed that the macrophages from obese mice are more infected by the L. major and it is worst in the presence of IL-17. Our results suggest that diet induced obesity decrease the resistance to infection.

## Introduction

Obesity is characterized by excessive fat accumulation, and it is considered a multifactorial chronic disease that has increased globally in westernized world over the last decades. It is associated with metabolic syndrome that includes insulin resistance, type 2 diabetes mellitus, dyslipidemia and hypertension, and also leads to respiratory diseases, hepatic steatosis, polycystic ovary syndrome, infertility, cancer, stroke, osteoarthritis, among others (1).

Metabolic syndrome and co-morbidities associated with obesity occur in an environment characterized by the presence of a chronic low-grade inflammation (2,3). The link between obesity and inflammation started to be established in early 1990’s when researchers demonstrated that TNF-α expression was elevated in adipose tissue and that it was related to insulin resistance (4). Since then, many studies have confirmed that adipose tissue produces cytokines with pro-inflammatory characteristics (5) that are responsible for increased macrophage recruitment (6,7) and for decrease in dendritic cell (DC) and natural killer cell (NK) functions (8,9). In addition, obesity alters the profile of T cells in adipose tissue (10). While T helper 1 (Th1) and T CD8+ are increased in the adipose tissue of obese animals (7), regulatory T cells are reduced (11,12). These alterations in immune cells may cause alterations in immune responses in obese individuals.

It was described that obesity increases the susceptibility to infection by different agents such as influenza virus (H1N1) (13), *Helicobacter pylori* (14) and *Staphylococcus aureus* (15). Karlsson and coworkers showed that obese mice infected with H1N1 virus had increased production of TNF-α and IL-6. Nevertheless, these animals had a poor memory TCD8+ cell response and were more susceptible to infection (16). Overweight and obese individuals also have a defective immune response to H1N1 viral infection (17).

Other studies have addressed the effect of obesity on parasite infections. There is a positive correlation between obesity and increased incidence of *Toxoplasma gondii* infection (18).

Interestingly, diet-induced obesity in C57BL/6 mice was protective in a model of cerebral malaria (19). Hypothalamic obesity in C57BL/6 mice infected with *Plasmodium berghei* ANKA resulted in decreased parasitemia, but exacerbated inflammation, and increased mortality rate (20). Leptin-deficient obese mice (ob/ob mice) are also more susceptible to *Trypanosoma cruzi* infection (21,22). Sarnáglia and coworkeres showed that diet-induced obesity promoted susceptibility to visceral leishmaniasis followed by higher production of pro-inflammatory cytokines and increased parasite load (23).

Together, these results describe contradictory phenomena: obesity causes increase in inflammatory immune response, but the increased inflammatory response did not lead to effective control of the microorganisms nor disease.

Resistance to *Leishmania major* infection is well characterized in C57BL/6 mice. Induction of an early Th1 response is necessary to induce resistance. Initial activation of dendritic cells leads to production of IL-12 (24) that promotes a Th1 response with high levels of IFN-γ and TNF-α production, and low levels of IgG1 antibody secretion. The establishment of this polarized inflammatory environment activates expression of iNOS and NO production in macrophages, which has leishmanicidal activity (25,26). Since this is a well established model of resistance to infection, we decided to investigate if obesity would interfere in the outcome of cutaneous leishmaniasis in C57BL/6 mice.

In this study, we showed that obesity did not affect the development of a Th1 response, nor triggered a Th2 or regulatory immune responses. However, obese mice were more susceptible to infection. Moreover, the increased IL-17 production found in obese mice in response to infection was not able to control leishmania growth *in vitro*, suggesting that this cytokine may favor parasite growth. The observed augment of IL-17 response against the parasite provides a mechanism for the increased susceptibility of obese mice to *L. major* infection. Our results can bring new insights into the relationship between immune response and obesity in animals facing a parasite infection with possible repercussions to a clinical scenario of leishmaniosis in obese individuals.

## Methods

### Animals and diet-induced obesity

All experiments were performed using six-eight-week-old female C57BL/6 mice, weighting approximately 18g, and obtained from the Animal Facility at the Universidade Federal de Minas Gerais (CEBIO, UFMG – Belo Horizonte, Brazil). All animals were maintained in the Experimental Animal Facility of Laboratório de Imunobiologia in collective cages (5 animals/cage) in an environmentally controlled room with a 12-hour light/dark cycle, controlled temperature (28°C) and unlimited access to water and food. Procedures and manipulation of animals followed the guidelines of the committees of ethics in research of Universidade Federal de Minas Gerais in agreement with guidelines of the committees of ethics in research according with Federal Law #11794, October 8th 2008: http://www.planalto.gov.br/ccivil_03/_ato2007-2010/2008/lei/l11794.htm.

All animal protocols were approved by the Committee on Animal Experiments (CETEA) under the protocol 338/2012, and it was approved in 01/10/2013. This certificate expires in 01/10/2018.

The obesity was induced with high sugar and butter diet (HSB), given *ad libidun* to the mice (12). The chow was full in micronutrients to do not induce nutritional deficit in mice. Experimental diet and the control diet was developed according published before by Reeves, 1993 (27).

### Experimental design

Mice were divided into two groups: control (fed AIN-93G) and obese (fed HSB). Animals were fed the same diet throughout the experiment. On the 4^th^ week of diet consumption, mice were infected with 1×10^6^ metacyclic promastigotes of *Leishmania major.* Infection was followed for 8 weeks; body weight, fasting glycaemia, and LDL cholesterol and triglycerides levels were also measured during this periods. Euthanasia occurred on the second, fourth and eighth weeks post infection (Fig 1A).

**Fig 1:**
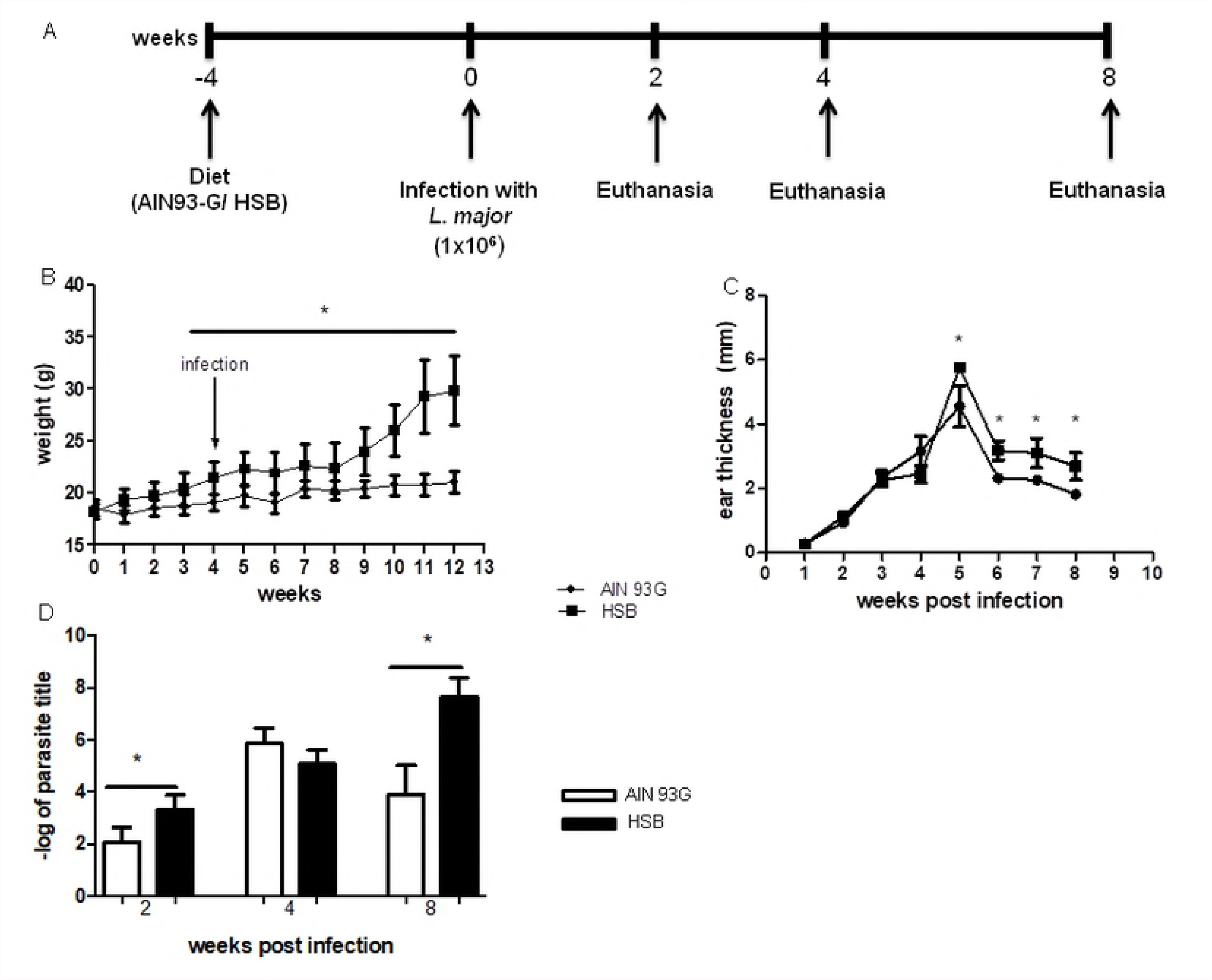
Experimental design and course of infection in diet-induced obese C57BL/6 mice. Animals were fed control (AIN93-G) or hypercaloric (HSB) diets *ad libitum* for four weeks. On the fourth week mice were infected with 1×10^6^ metacyclic *L. major* in the ear. Weight gain and food intake were measured weekly. (A) Experimental design. (B) Weekly measurement of C57BL/6 body weight. (C) Weekly measurement of infected ear thickness during eight weeks of infection. (D) Parasite titer in the infected ear, measured by limiting dilution. Statistical analysis was performed by Student’s *t* test (*p<0.05; **p<0.005). Results are representative of at least 4 independent experiments, n=4 mice/group.

### Parasites, infection and antigens

*Leishmania (Viannia) major* (WHOMHOM/IL/80/Friedlin) were maintained in Grace‘s medium (GIBCOBRL – Life Technologies, Grand Island, NY, MO, EUA), pH 6.2 supplemented with 20% of fetal bovine serum (GIBCO), 20μg/mL gentamicin sulfate (Schering-Plough – Rio de Janeiro, RJ, Brazil) and 2mM de L-glutamine (GIBCOBRL – Life Technologies, Grand Island, NY, MO, EUA) (supplemented Grace‘s), at 25°C. For infection metacyclic promastigote were used 1×10^6^, obtained after 5 days in culture. The parasites were inoculated intradermically in the left ear of each animal (final volume = 10μl). Lesion development was monitored weekly by the difference in thickness between infected and uninfected ears.

Number of parasites was estimated by limiting dilution as described previously (28). Briefly, mice were euthanized and the whole ear was removed and cleaned in 70% alcohol. Ears were fragmented with scissors and grinded in a glass tissue grinder. Tissue debris were removed by centrifugation at 50 *g* for 1 min and the supernatant was transferred to another tube and centrifuged at 1,540 *g* for 15 min. Pellet was resuspended in 0.5ml supplemented Grace’s medium. The parasite suspension was then serially diluted in 10-fold dilutions in duplicates to a final volume of 200µl in 96-well plates. Pipette tips were replaced for each dilution. Plates were incubated for 10 days at 25°C and examined under an inverted microscope. Results were expressed as the negative log of the last dilution in which parasites were detected.

*Leishmania* antigen was obtained from logarithmic phase cultures of *L. major* promastigotes. Promastigotes were washed twice in PBS and pellets were submitted to seven cycles of freezing in liquid nitrogen followed by thawing at 37°C. The preparations were visually inspected for the presence of intact parasites. Protein content of preparations was assayed by the Lowry method (29) and adjusted to 1mg/ml protein. Antigen preparation was aliquoted and stored frozen at - 20 °C.

### Histology

After euthanasia, ear samples were collected and fixed in 80% methanol and 20% dimethyl sulfoxide (DMSO; Merck, Darmstadt, Germany), embedded in paraffin, cut into 3–5-μm sections and stained with hematoxylin–eosin (H&E-staining) for microscopic analysis. Images were taken in optical microscopic and are shown in 100x augmentation.

### Glycaemia and LDL measuremets

To perform the glycemic ratings, animals were fasted for 6 hours and blood was collected from the tail vein. Blood glucose was measured in with a glucometer and strips (Accu - Chek Performa^®^). To perform the oral glucose tolerance test (OGTT), glucose was given by gavage at a concentration of 2g/kg body weight. Measurements were performed with a glucometer and strips (Accu - Chek Performa^®^) before glucose gavage and at 15, 30, 60 and 90 minutes later. For fasting glucose and LDL cholesterol measurements animals were fasted for 6 hours and blood was collected. The glycaemia and LDL cholesterol levels were measured by enzymatic kit according to fabricant‘s protocol (Bioclin, Quiabasa, MG, Brazil).

### Enzyme-linked immunosorbent assay (ELISA) for cytokines and antibodies detection

For cytokines measurement, plates were coated with 50μL/well with monoclonal antibodies solution (against IFN- γ, TNF- α, IL −4, IL −10 and IL −17 (1μg/ml) (PeproTech, NJ, US), diluted in PBS and kept overnight, at room temperature. The enzymatic reaction was revealed by incubating the plates with a solution containing 0.2 μL/mL of H_2_O_2_ and 0.5 mg/ml ABTS ((C_18_H_16_N_4_O_6_S_4_-(NH_4_)_2_) (Sigma-Merk, Germany) in xM citrate buffer pH 5.0 for the development of a dark green coloration. After this stage, the reactions were stopped by addition of 20μL/well of a solution of SDS 1%. The absorbance (λ405nm) of each well was obtained by automatic ELISA reader (Molecular Devices Spectra MAX340).

### Macrophage culture

Thioglycolate 2% solution was inoculated in the animal’s peritoneum, at 8 weeks after diet consumption, to induce macrophage recruitment. Peritoneal exudate cells were obtained after 72 hours. The collected macrophages were incubated in a 24 well plate onto glass coverslips at the concentration of 1×10^5^ cells/ml. Cells were incubated for 2 hours for adhesion and received 5 *L. major* in the stationary phase per macrophage. *In vitro* infection was analyzed by optical microscopy 4, and 72 hours post infection after instant glass coverslip staining (Panótico, Laborclin, PA, Brazil) For nitric oxide (NO) and arginase activity, cells were incubated in 96 well plates in the concentration of 1×10^6^ cells/ml and infected with 5:1 *L. major*. The supernatant was collected for nitric oxide measurement at 72h post infection, as well as arginase activity. Cells were stimulated wiht 1ng/ml of IFN-γ, 1ng/ml of IL-4 and 1ng/ml of IL-17, and 100ng/ml of LPS.

### Nitric oxide (NO) detection

Supernatant of macrophage cultures were collected 72h post *in vitro* infection. Nitric oxide production was measured as nitrite in culture supernatants using the Griess’ reaction (30).

### Arginase Activity

Arginase activity in homogenates of infected macrophages was assayed as described previously (31) with few modifications. About 35μL macrophage homogenate was incubated in 24-well plates with 50μL Triton X-100 and plates were shaken for 30min. Arginase was activated with 50μL MnCl_2_ (10mM) and 50μL of TRIS-HCl (50mM, pH 7.5) at 55°C for 10 min. Then, 50μL samples were transferred to a fresh 24-well plate containing 25μL of L-arginine (0.5mM, pH 9.7) and incubated for 60 min at 37°C. The reaction was stopped by the addition of 400μL of a mixture of acids and water H_2_SO_4_:H_3_PO_4_:H_2_O (1:3:7). Subsequently, 25μL of 9% 1-phenyl-1,2- propanodione-2-oxime in ethanol was added and the plates were incubated at 95°C for 45 min for color development. Reaction mixtures were read at 540nm in a spectrophotometer (Molecular Devices). One unit of enzyme activity is defined as the amount of enzyme that catalyzes the formation of 1μmol of urea/min. A standard curve was performed using urea and the detection limit for the assay was 270μM of urea.

### Statistical analysis of the data

Data were initially analyzed using the Kolmogorov-Smirnov test to verify normal distribution. Since all data were normally distributed, Student’s *t* test and one-way ANOVA were used to compare groups. The significance level of p<0.05 was adopted.

## Results

### Obesity was associated with higher parasite burden

C57BL/6 mice fed HSB-diet started to develop obesity in the 4^th^ experimental week. Obese mice presented a significant increase in body weight when compared with the control group (Fig 1B). Obesity persisted until the end of the study. We also confirmed the prevalence of metabolic syndrome in HSB-fed mice by their high levels of blood glucose, altered glucose tolerance test, high serum levels of LDL cholesterol, triglycerides, and leptin (Sup Fig 1).

In order to verify whether obesity would affect the course of infection with *L. major* in C57BL/6, mice were infected in the left ear with metacyclic promastigotes 4 weeks after HSB-diet consumption. Ear thicknesses in obese mice were significantly higher than in control mice from the fifth week post infection on (Fig 1C). Parasite burden was also checked 2, 4 and 8 weeks post infection. Two weeks post infection and eight weeks post infection obese mice presented higher parasitism (Fig 1D). Histological analysis of ear samples were performed to search for differences in the lesion inflammatory profile, and animals from obese group had more parasites in the second week post infection. In the eighth week post infection, obese mice presented larger cellular infiltrate, including polymorphonuclear cells and mastocyte hyperplasia (Fig 2D, E and F).

**Fig 2:**
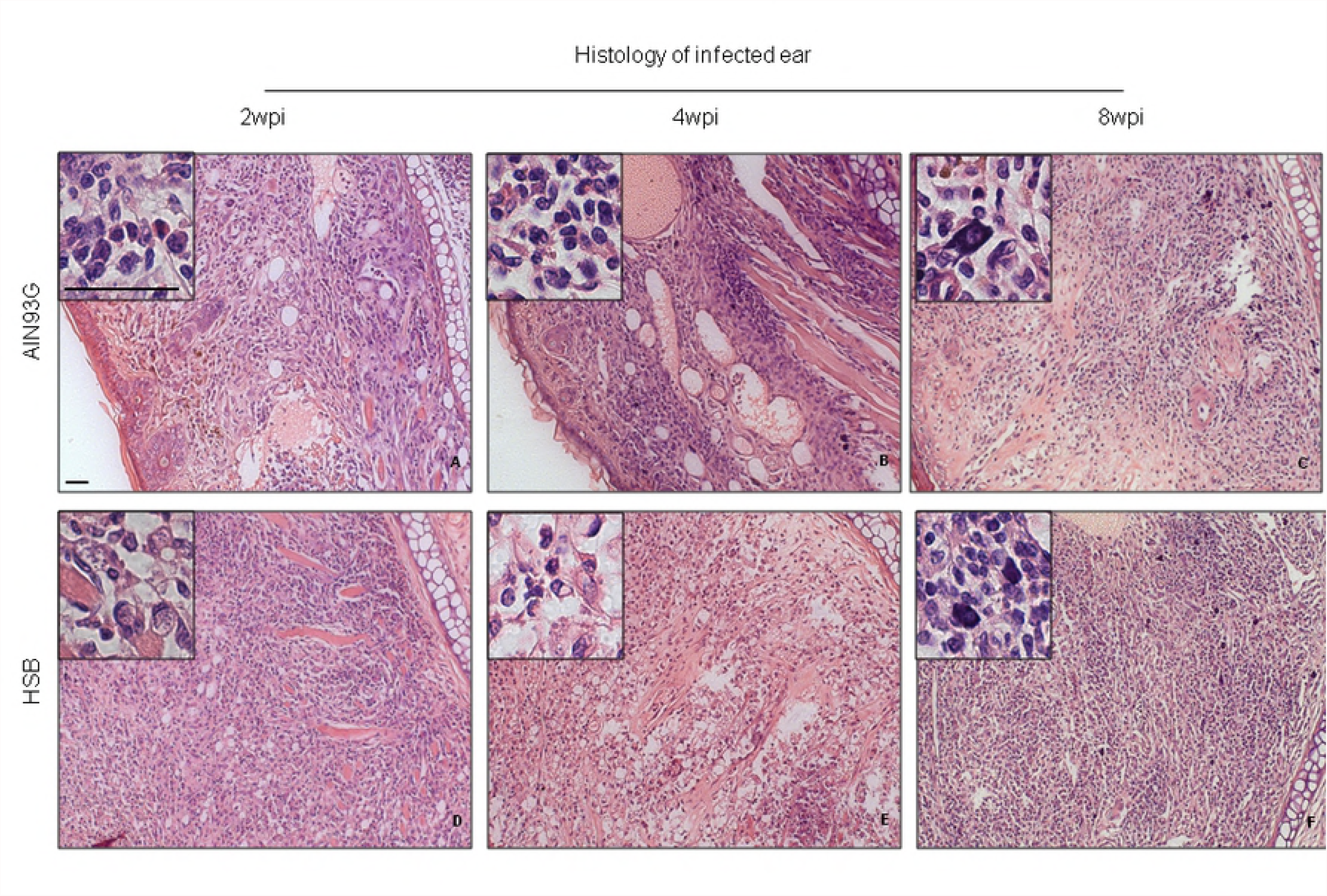
HSB diet-induced obesity leads to more inflammation in the infected ear. Representative photomicrographs of ear histological sections from C57BL/6 mice submitted to HSB and AIN93-G diets and infected with *L. major*. (A, B and C) photomicrography from control mice 2, 4 and 8 weeks post infection. The cellular infiltration is predominantly polymorphonuclear (Insert in A, B), focal and discrete to moderate infiltration in the papillary dermis and deep dermis accompanied by mild to moderate thickening of the dermis; erosion and ulceration of the epidermis, predominantly polymorphonuclear inflammatory process with discrete mast cell hyperplasia (Insert in C) and thickening of the dermis, moderate to severe at eight weeks post infection. (D, E and F) photomicrography from HSB-fed mice 2, 4 and 8 weeks post infection. Ulcerated epidermis, predominantly polymorphonuclear and focal inflammatory process in the dermis and hypodermis, with thickening of the dermis, both of intense character, and tissue parasitism (Insert in D) in animals belonging to the obese group, 2 weeks post infection; moderate thickening of the dermis and inflammatory infiltrate also moderate, predominantly polymorphonuclear (Insert in E) and focal, in the papillary dermis, deep dermis and hypodermis 4 weeks post infection; ulcerated epidermis, predominantly polymorphonuclear and focal inflammatory process in the dermis and hypodermis with thickening of the dermis, both moderately characterized with moderate mast cell hyperplasia (Insert in F), 8 weeks after infection. Hematoxylin-Eosin. Bar = 25mm.

### Obesity induced increased production of *L. major* specific IgG1

In the eighth week post infection, obese mice presented higher levels of specific circulating IgG, IgG1 and IgG2a, when compared to the levels found in mice from the control group (Fig 3A, C and D). Serum IgM levels were higher in obese mice only on the 4^th^ week. There was an increase in total IgG, IgG1 and IgG2a levels on the eighth week of infection in obese mice when compared to their control counterparts suggesting that increased antibody production during obesity may be associated with a lower efficiency to kill *Leishmania*.

**Fig 3:**
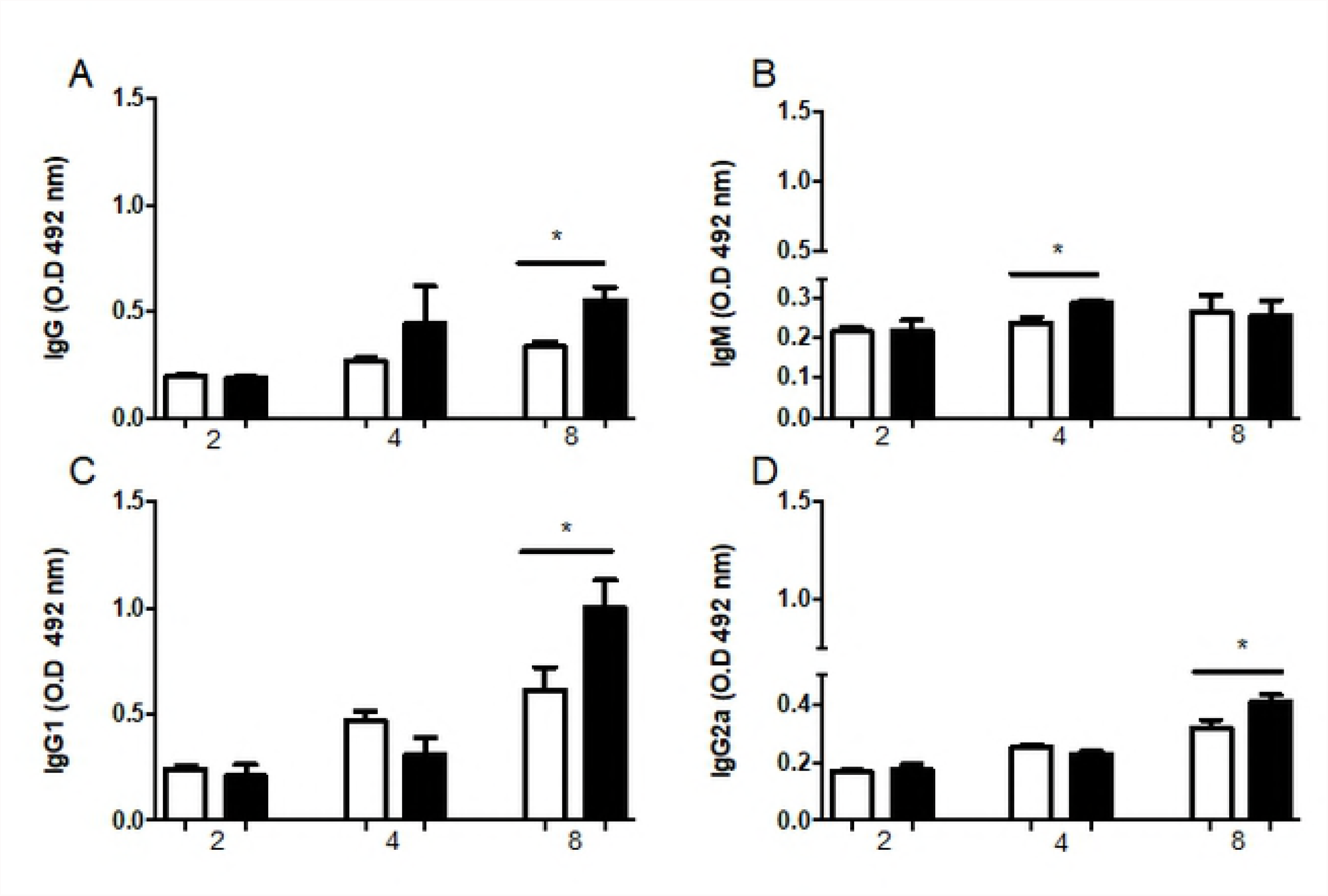
Diet-induced obesity increases the concentration of *L. major* specific immunoglobulins in C57BL/6 serum. Total IgG (A), IgM (B), IgG1 (C) and IgG2a (D) were measured in sera from mice fed control (AIN93G) and hypercaloric (HSB) diet. ELISA was performed at 2, 4 and 8 weeks of infection. For sensitization, *L. major* antigen at 20µg/ml was used. To measure IgG, sera were diluted 1:1000, and 1:100 for IgG1, IgG2a and IgM. Statistical analysis was performed by Student’s *t* test (\**p<0.05*; \*\**p<0.005*). Results are representative of at least two independently experiments, n= 4 mice/group.

### Obese C57BL/6 mice showed no impairment in IFN-γ production, but had increased IL-17 production by draining lymph node cells

Infection with *L. major* in C57BL/6 mice induces a typical Th1 immune response with production of high levels of IFN-g that activates macrophages, a cell type directly responsible for parasite control (Hurdayal and Brombacher, 2017). Both control and obese mice produced equivalent levels of IFN-γ by cells from the draining lymph node (Fig 4A). The same result was observed for TNF-α (Fig 4B). These data suggest that the increase in parasitism observed in obese mice was not related to a deficient Th1 response.

**Fig 4:**
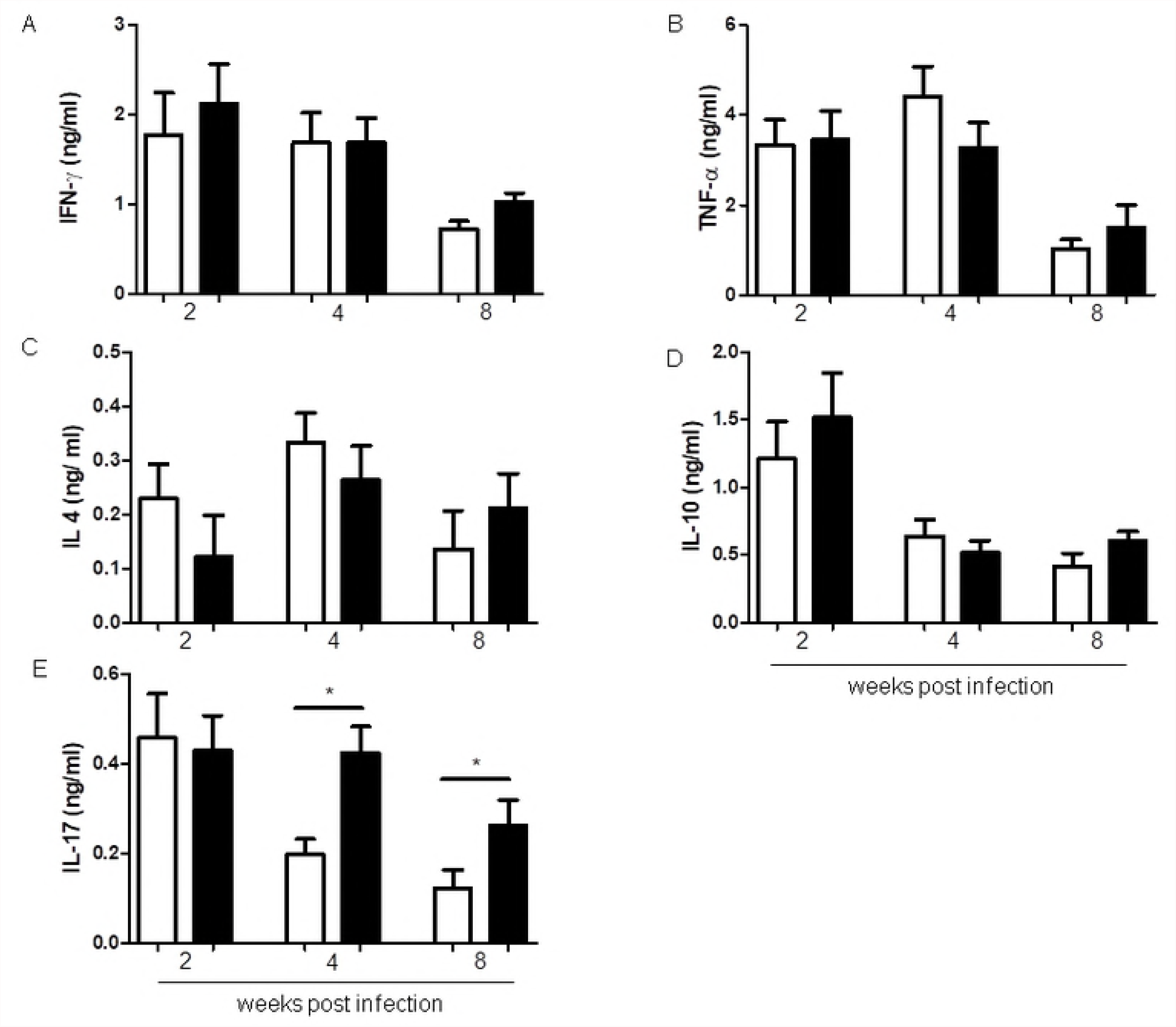
Cytokine production by lymph node cells in culture. Cytokine concentrations were measured by ELISA in cell culture supernatants stimulated *in vitro* with 50µg/ml of *L. major* antigen. Cells were collected 2, 4 and 8 weeks after infection and adjusted for 5×10^6^/mL of culture and incubated during 72h. (A) IFN-γ; (B) TNF-α; (C) IL-4; (D) IL-10 and (E) IL-17. Statistical analysis was performed by Student’s *t* test (* p<0.05). Results are representative of at least two independent experiments, n=4 mice/group.

We also evaluated the production of IL-4, IL-10 and IL-17 in cultured cells from draining lymph nodes. There was no difference in IL-4 or IL-10 secretion between obese mice and control mice (Fig 4C and 4D). Interestingly, IL-17 production was high on the second week post infection in both groups. However, in obese mice IL-17 levels stayed high up to 8 weeks of infection, while they decreased in control mice. IL-17 production has been associated with obesity (32), and also with uncontrolled cutaneous *Leishmania* infection (33).

Cytokine secretion in adipose tissue was also evaluated and no difference was found for IFN-γ nor for IL-17. As expected, TNF-α, levels were increased in adipose tissue from obese mice before infection, they were decreased two weeks post infection, and there was no difference between control and obese mice four and eight weeks post infection. IL-10 production in adipose tissue also presented some differences between obese and control mice. In obese mice, IL-10 levels were higher before infection and four weeks after infection (Supplementary Fig 2).

### Macrophages from obese mice had higher arginase activity and higher parasitism when infected *in vitro* with *L. major*

To evaluate macrophage activity in obese mice, we analyzed the degree of infection of macrophages at times 4 and 72 hours post *in vitro* infection with *L. major*. At 4 hours post infection, macrophages from obese mice had a higher percentage of infected macrophages as well as a higher number of amastigotes per macrophage when compared to macrophages from control mice with obesity (Fig 5A and B), and the infection index was also higher for macrophages from obese mice 4 hours post infection. Of note, *in vitro* infection was performed using total promastigotes, and that why there were less amastigotes 72h after infection than 4h post infection (34). At 72 hours post infection, macrophages from obese mice continued to harbor more amastigotes/cell than macrophages from control mice (Fig 5B). We observed that obese mice had their levels of IL-17 increased. To verify whether IL-17 would impair the killing of amastigotes *in vitro*, we stimulated the cultures with IL-17. It was observed that macrophages from control mice, when cultured *in vitro* with IL-17, had increased of amastigotes/cell and higher infection index 4 hours post infection (Fig 5B and 5C). At 72h post infection, macrophages from control mice controlled the infection, especially when stimulated with IFN-γ. Moreover, addition of IL-17 to the cultures impaired the clearance of amastigotes in infected cells as seen by the infection index in both control and obese groups. These data suggest that IL-17 may interfere with parasite elimination.

**Fig 5:**
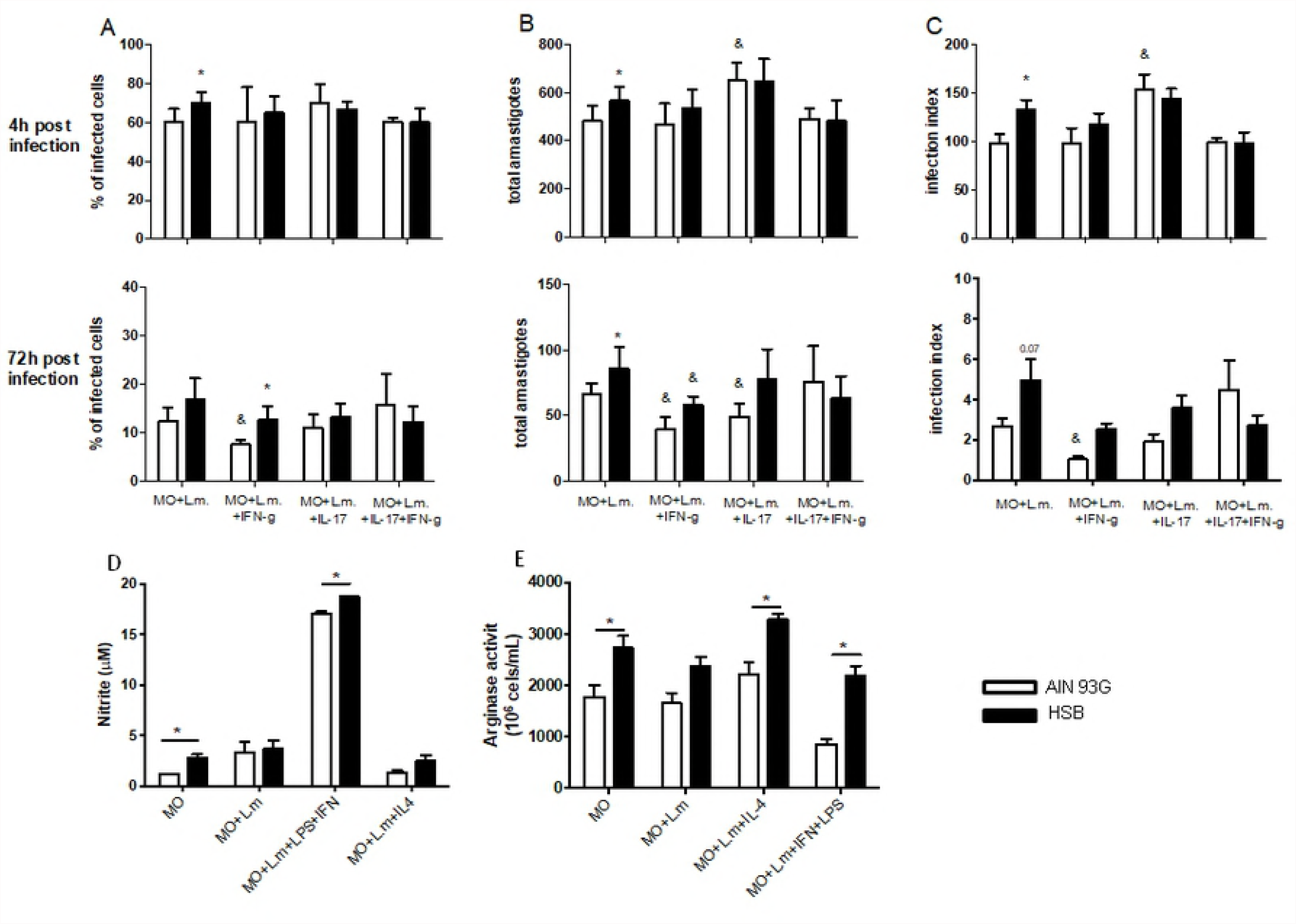
*In vitro* macrophage infection. To measure the percentage of infected macrophage and the amastigote number in infected macrophage, cells were collected from the peritoneal cavity after thioglycotate stimulation and cultured at 1×10^5^/ml. The infections was performed with 5 parasites per macrophage 24h after adhesion. Parasite counts were performed at 4 and 72 hours post infection. Arginase and NO were measured 72h after infection, as described in Materials and Methods. (A) Percentage of infected macrophages; (B) Amastigote number per infected macrophage and (C) Infection index; (D) NO production; and (E) Arginase activity. Statistical analysis was performed by Student’s *t* test (*p<0.05 between macrophages from control mice versus obese mice; ^&^ p<0.05 comparing different culture conditions in the same group). Results are representative of two independently experiments, n=3).

We also investigated the production of NO and arginase activity by macrophages from obese mice. Macrophages from obese mice without any stimulus or stimulated with IFN-γ and LPS produced slightly more NO than cells from control group (Fig 5D). However, arginase activity was also higher in non-stimulated cells from obese mice, as well as in cells infected with *L. major.* IL-4 stimulation did not increase arginase activity, but arginase activity remained higher in obese mice. Macrophages from obese mice also displayed higher arginase activity upon activation with IFN-γ and LPS (Fig 5E). Therefore, our data suggest that obesity increases arginase activity in macrophages.

## Discussion

The impact of obesity in the immune response to cutaneous leishmaniasis has never been described before. Our studies showed that C57BL/6 mice with diet-induced obesity present larger lesions than control mice, and these lesions contained more parasites. We further investigated the mechanisms that could be causing these differences.

There is a gap in the understanding of how obesity affects the course of intracellular infections, including parasite infections. In accordance with the negative association between obesity and infectious diseases already described (23,35,36), our study showed that obese mice had higher parasite burden than control mice, failing to control parasite growth. In addition, lesions in obese mice were larger and more ulcerative than the ones observed in the control group. Adipose tissue produces cytokines, adipokines, and chemokines which alter the immune response affecting the recruitment of inflammatory cells such as macrophages, neutrophils and dendritic cells (37). Obesity also affects T cell differentiation and function, increasing pro-inflammatory cytokine production (5,38). Considering the “low-grade” inflammation seen in obesity, one reasonable hypothesis could be that, in the specific case of cutaneous leishmaniasis, where macrophages require an inflammatory environment to control parasites, obesity would improve the immune response against *L. major*. However, our data do not support this hypothesis.

Our histological data also detected a higher parasite burden in obese C57BL/6 mice when compared to mice from the control group further indicating that obesity interferes with control of parasite growth. Interestingly, lesions of obese mice presented higher degree of cellular infiltration, but the inflammatory cells had a poor ability to eliminate the parasites.

We also measured the antibody response to *L. major* antibody in sera, and found higher levels of IgG and IgG1 in obese C57BL/6 mice than in control mice eight weeks post infection. As described in previous studies, susceptibility to *L. major* infection is associated with isotype switching to IgG1 (39,40). Antibody levels are directly related to parasite number, as antibodies may form immune complexes that bind to Fcγ receptors (FcγR), inhibiting the proinflammatory activity of macrophages, without impairing phagocytosis (41). Interestingly, despite favoring phagocytosis, IgG fails to protect against *L. major*, and even worse, contributes to the pathogenicity itself (42). Indeed, it has been reported that phagocytosis of IgG-opsonized amastigote forms induces the activation of signaling pathways leading to the production of IL-10 by macrophages (40).

In an attempt to understand why obese C57BL/6 mice had more severe lesions than control mice, we evaluated cytokine production by these mice. Interestingly, obesity did not affect the production of IL-4 in C57BL/6 mice. This result differs from previous data that associate obesity with susceptibility to asthma and allergies, driven by higher IL-4 secretion and Th2 differentiation (43,44). In fact, the balance between Th1 and Th2, as well the cytokine profile during obesity are controversial. We also showed that TNF-α and IFN-γ levels were increased in both groups at the time points measured. Obese mice did not show overproduction of proinflammatory cytokines by cells from the draining lymph nodes. It was expected a higher production of TNF-α and IFN-γ in obese mice with in response to parasite antigens, as it was observed for infection with *Plasmodiun berguei* (45) and also for infection with *Leishmania chagasi* (23) in obese C57BL/6 mice. However, obesity did not interfere with TNF-α and IFN-γ levels in our study.

The role of IL-17 in leishmaniasis has been a subject of debate. Moreover, the role of IL-17, in general, is still controversial. We found a higher IL-17 production in obese mice, which presented a more severe lesion with higher parasite burden than the control group. Classically, IL-17 production is associated with neutrophil recruitment and often is related to proinflammatory response in various diseases, including autoimmune disorders (46), fungal (47)) and bacterial infections (48). However, recent studies have questioned whether IL-17 function is restricted to proinflammatory action (49). IL-17 is produced mainly by Th17 cells, which could be stimulated by a different combination of cytokines, including TGF-β, IL-6, IL-23 and IL-1β (46). Obesity alters the cytokine profile in adipose tissue and in serum, and IL-17 seems to have a significant role in obesity. Previous studies have shown that production of this cytokine is elevated in obesity, both in humans and mice (32,50). In line with these reports, our results showed that cells from drainig lymph node of obese infected mice secrete higher levels of IL-17. Studies on cutaneous leishmaniasis have correlated IL-17 release with increased pathogenicity in cutaneous leishmaniasis in mice (51,52). Therefore, it is plausible that the increased susceptibility of obese C57BL/6 to Leishmania infection in our model is related to increased IL-17 production by draining lymph node cells.

We also performed *in vitro* infection of macrophages with *L. major* to assess the efficiency of macrophages from obese C57BL/6 mice to kill the parasites. There was an increased frequency of infected cells among macrophages from obese mice 4 hours post infection, and a larger number of amastigotes/cell in macrophages from obese mice 4 and 72 hours post infection. These results are in line with higher arginase activity detected in macrophages from obese mice. Interestingly, Sousa and coworkers found that IL-17 increases arginase activity, and favor parasite growth in BALB/c mice infected with *L. amazonensis* (33). The hallmark of a Th2 response is activation of M2 macrophages and induction of arginase 1, which requires IL-4 produced by Th2 cells. Arginase activity is increased in mice susceptible to cutaneous leishmanisis, and this enzyme utilizes arginine as a substrate to induce polyamine production instead of NO (53). Interestingly, studies on inflammatory bowel disease have demonstrated that IL-17 induces a “M2-like response”. Moreover, IL-17KO C57BL/6 mice expressed lower levels of mRNA coding for molecules associated with M2 activity, including arginase 1 (54). Another study showed that IL-17A induces arginase 1 production in a model of human Papillomavirus (55). In this case, arginase 1 would be active in M2-macrophages. Our *in vitro* results showed an environment where IFN-γ would still be inducing an inflammatory response in obese mice. In spite of that, IL-17 production compromised the leishmanicidal activity of macrophages from obese mice and further facilitated parasite growth by increasing arginase activity.

To test the hypothesis that IL-17 could impair leishmania killing by macrophages, we infected macrophages stimulated *in vitro* with IL-17. IL-17 had no effect on macrophages from obese mice. However, IL-17 decreased parasite killing by control macrophages activated with IFN-γ.

Classically, a Th1 response would lead to resistance to *L. major,* while a Th2 response would lead to susceptibility (56). However, this classical paradigm has been challenged by later studies on *L. major* infection (39,57). In the present work, we propose IL-17 as an alternative cytokine that may determine the fate of the immune response against *L. major*. We showed that diet-induced obesity, a condition associated with IL-17 production, increased susceptibility of C57BL/6 mice, a mouse strain genetically resistant to *L. major* infection. Therefore, IL-17 could be a potential candidate to explain diverse comorbidities associated with obesity and its role in models of infection should be better explored.

Taken together, our results show that diet-induced obesity in C57BL/6 mice decreased their capacity to control infection by *L. major*. This might be related induction of IgG1 secretion, IL-17 production and impaired capacity of macrophages to control parasite growth. The present provides novel clues to the relationship between obesity and leishmanial infection in a time when infection is now being added to the list of health risks associated with obesity.

## Acknowledgments

We would like to thank Ilda Marçal and Hermes dos Reis Ribeiro for her excellent care of the animal facility, and Maria do Carmo Dias for technical help. We thank Professor Simone Vasconcelos Generoso for providing the recombinant cytokines used.

## Figures Legends

**Sup Fig 1: Metabolic evaluation in mice fed HSB diet before and after *L. major* infection.** (A) Fasting glycaemia. (B) Glucose oral tolerance test performed eight weeks post infection. Mice were fasted for six hours and received a 30% glucose solution *per os.* Blood was taken at time 0 and 15, 30, 60 and 90 minutes after administration of the glucose solution. (C) Total blood cholesterol eight weeks post infection. (D) Serum leptin concentration measured by ELISA (8 weeks post infection and 12 weeks post diet consumption). Statistical analysis was performed by Student’s *t* test (* p<0.05; ** p<0.005 and *** p<0.0005). Results are representative of at least 4 independently experiments, n=4mice/group.

**Sup Fig 2: Cytokine profile in the adipose tissue extract from C57BL/6 mice infected with *L. major*.** The peritoneal adipose tissue extracts were prepared (100mg/ml of buffer) and ELISA was performed to measure concentrations of IFN- γ, TNF-α, IL-10, IL-17 and IL-4. (A) IFN- γ; (B) TNF-α; (C) IL-10; (D) IL-17. IL-4 values were below the detection limit. Statistical analysis was performed by Student’s *t* test (* *p*<0,05). Results are representative of at least two independently experiments, n=4mice/group.

